# Activation of SARS-CoV-2 by trypsin-like proteases in the clinical specimens of patients with COVID-19

**DOI:** 10.1101/2022.09.27.509803

**Authors:** Emiko Igarashi, Takahisa Shimada, Shunsuke Yazawa, Yumiko Saga, Masae Itamochi, Noriko Inasaki, Sumiyo Hasegawa, Yoshitomo Morinaga, Kazunori Oishi, Hideki Tani

## Abstract

SARS-CoV-2 enters host cells through the angiotensin converting enzyme 2 (ACE2) receptor and/or transmembrane protease, serine 2 (TMPRSS2). Serine proteases, such as TMPRSS2 and trypsin, promote viral entry.

In this study, we investigated whether proteases increased SARS-CoV-2 infectivity using pseudotyped viruses and clinical specimens from patients with COVID-19. First, we investigated how trypsin increased infectivity using the pseudotyped virus. Our findings revealed that trypsin increased infectivity after the virus was adsorbed on the cells, but no increase in infectivity was observed when the virus was treated with trypsin. We examined the effect of trypsin on SARS-CoV-2 infection in clinical specimens and found that the infectivity of the SARS-CoV-2 delta variant increased 36,000-fold after trypsin treatment. By contrast, the infectivity of SARS-CoV-2 omicron variant increased to less than 20-fold in the clinical specimens. Finally, infectivity of clinical specimens containing culture supernatants of *Fusobacterium necrophorum* was increased from several- to 10-fold. Because SARS-CoV-2 infectivity increases in the oral cavity, which may contain anaerobic bacteria, keeping the oral cavities clean may help prevent SARS-CoV-2 infection.

**Importance:** In this study, we examined whether trypsin-like proteases increased the infectivity of SARS-CoV-2. We found that trypsin-like proteases increased the infectivity of both the pseudotyped viruses and the live virus in the clinical specimens. The increase in infectivity was significantly higher for the delta than the omicron variant. A large amount of protease in the oral cavity during SARS-CoV-2 infection is expected to increase infectivity. Therefore, keeping the oral cavity clean is important for preventing infection.

## Introduction

At the end of 2019, the outbreak of coronavirus disease 2019 (COVID-19) occurred in China and soon turned into a global pandemic, with more than 600 million people infected and nearly 6.5 million dead as of September 2022. Subsequently, several variants of SARS-CoV-2 have emerged, and various countermeasures are being taken to prevent their spread. SARS-CoV-2 causes respiratory infections, and infects and proliferates in airway epithelial cells like typical influenza viruses. SARS-CoV-2 infects by endocytosis using the angiotensin converting enzyme 2 (ACE2) receptor; however, infectivity is enhanced by membrane fusion in the presence of the cellular serine protease, transmembrane protease, serine 2 (TMPRSS2) (1, 2).

The influenza virus is not infective unless hemagglutinin is cleaved by a protease. *In vivo*, the influenza virus becomes infectious when it is cleaved near airway epithelial cells by trypsin-like enzymes, such as tryptase Clara, which is released from Clara cells, or host cellular proteases present in the upper respiratory tract. In addition, studies have reported that staphylococcal proteases increase viral infectivity in the upper respiratory tract (3) and proteases from oral bacteria enhance SARS-CoV infection (4). Therefore, keeping the oral cavity clean and reducing the amount of proteases derived from indigenous bacteria may prevent infection. In fact, prevalence of influenza was found to be reduced when the oral cavity of residents was cleaned in elderly care facilities (5).

In this study, we examined whether SARS-CoV-2 is activated by trypsin-like proteases using clinical specimens from patients with COVID-19 and a SARS-CoV-2 surrogate model. Furthermore, we investigated whether proteases derived from oral anaerobic bacteria promote SARS-CoV-2 infection, and clarified whether cleaning of the nasal and oral cavities helps prevent SARS-CoV-2 infection.

## Result

### Trypsin treatment increases SARS-CoV-2pv infectivity

SARS-CoV-2 is highly regulated legally and must be handled in facilities with biosafety level (BSL) 3. Therefore, the Wuhan, alpha, delta, and omicron variants of SARS-CoV-2pv were generated using the vesicular stomatitis virus (VSV) pseudotyping system. Using these viruses allowed us to perform experiments in a BSL2 laboratory. In addition, because luciferase was inserted as a reporter gene, the efficiency of infection was quantitatively evaluated using luciferase activity as an index. Furthermore, the use of pseudotyped viruses eliminates the effects of components derived from clinical specimens.

Trypsin increased the infectivity of the Wuhan, alpha, and delta variants of SARS-CoV-2pv in a concentration-dependent manner—up to ∼10-fold at 800 μg/mL (Fig. 1A). By contrast, trypsin treatment did not increase the infectivity of the SARS-CoV-2pv omicron variant.

**Fig. 1.**
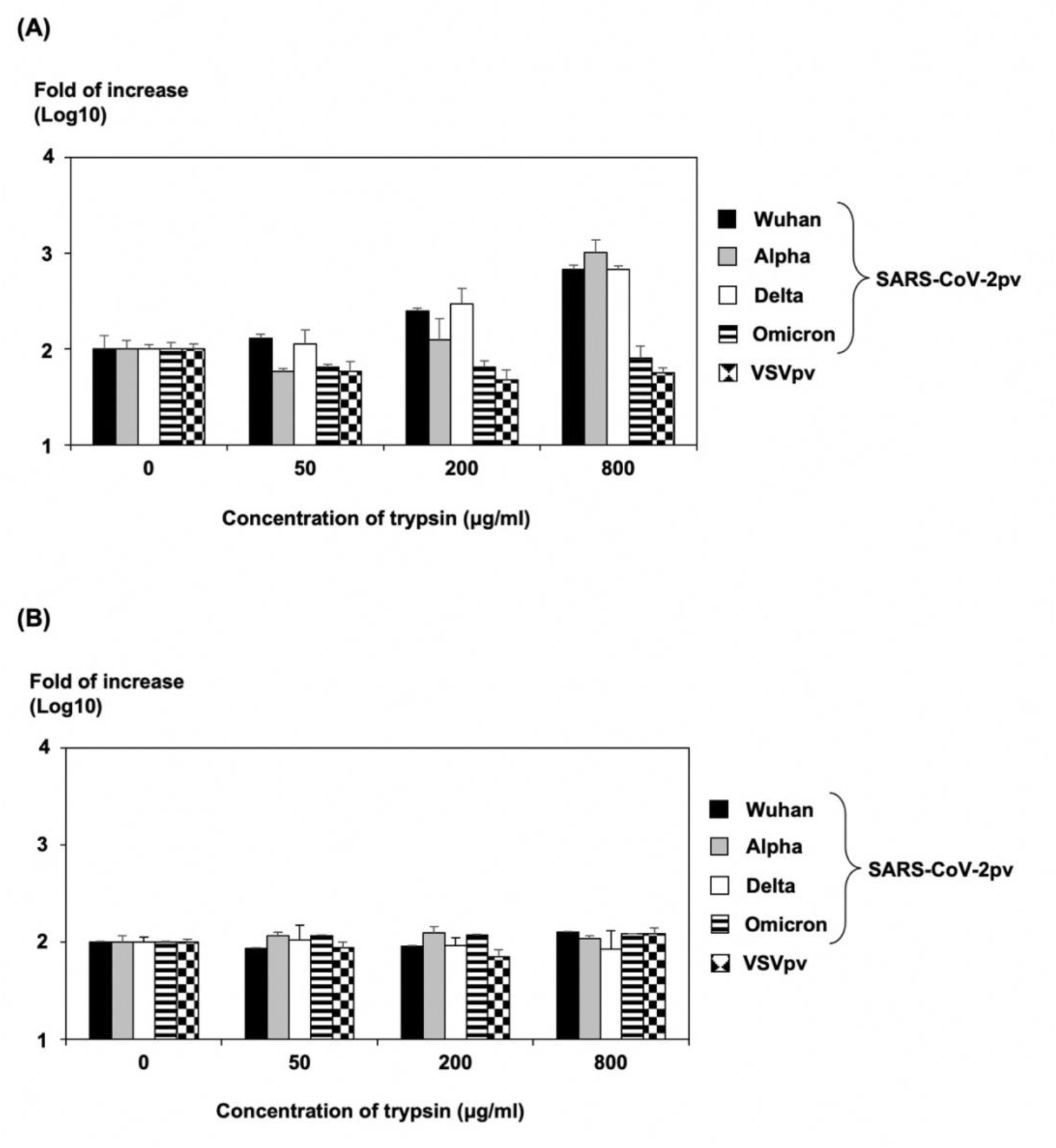
Effect of trypsin treatment on SARS-CoV-2pv infectivity. (A) The increased infectivity of viruses treated with trypsin at the indicated concentration after inoculation of cells with the Wuhan, alpha, delta, and omicron variants of SARS-CoV-2pv or VSVpv was calculated based on the infectivity of the trypsin-untreated virus. (B) The viral supernatants of Wuhan, alpha, delta, and omicron variants of SARS-CoV-2pv or VSVpv were inoculated into cells after treatment with trypsin at the indicated concentrations, and the increased infectivity of viruses was calculated based on the infectivity of the trypsin-untreated virus. The results shown are from three independent assays, with error bars representing standard deviations.

Next, we examined the action of trypsin treatment temporally and treated SARS-CoV-2pv with trypsin before infecting cells. We found no increase in infectivity effect in any variant (Fig. 1B). These results suggest that trypsin increases SARS-CoV-2pv infectivity after the virus is adsorbed on the cell and does not activate viral particles.

### Syncytium formation of S protein-expressing cells after trypsin treatment

To investigate whether the fusion of S protein of SARS-CoV-2 and host cellular membrane is involved in the trypsin-mediated increase in infectivity, we compared syncytium formation in cells expressing S protein after trypsin treatment (Fig. 2). VeroE6 cells expressing S protein of the Wuhan, alpha, and delta variants of SARS-CoV-2 formed particularly large syncytia after trypsin treatment (Fig. 2A). This phenomenon was confirmed in 293T cells expressing S protein of the SARS-CoV-2 delta variant, although cells expressing S protein of SARS-CoV-2 Wuhan and alpha variants exhibited weak fusion activity (Fig. 2B). Almost no syncytium formation was observed in cells expressing S protein of the SARS-CoV-2 omicron variant or VSVG in both cell types. These results indicate that the omicron variant shows weaker fusion activity than other variants after trypsin treatment.

**Fig. 2.**
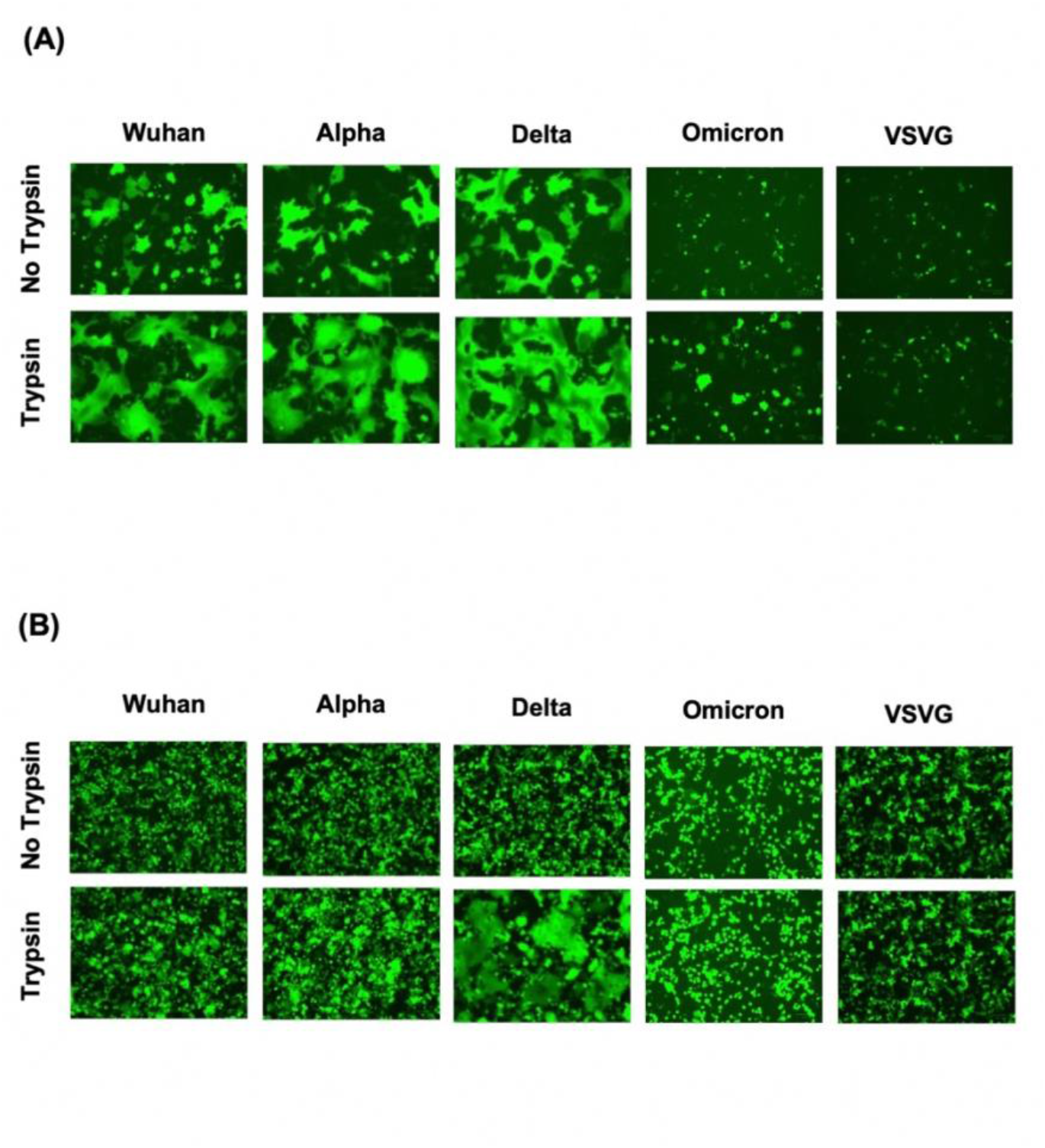
Effect of trypsin treatment on syncytium formation in cells expressing S protein of SARS-CoV-2. Syncytium formation in (A) VeroE6 and (B) 293T cells transiently expressing S protein of SARS-CoV-2. The eGFP-expressing plasmid was cotransfected for visualization. Syncytium formation was imaged using a fluorescence microscope.

### Trypsin treatment increases SARS-CoV-2 infection in clinical specimens

First, we examined the effect of trypsin treatment on SARS-CoV-2 infection in clinical specimens. We found that trypsin treatment increased the infectivity of 56 specimens containing the SARS-CoV-2 delta variant by 4.2- to 36,000-fold compared with untreated specimens (Fig. 3A). By contrast, the infectivity of 33 specimens among the 44 SARS-CoV-2 omicron variants increased after trypsin treatment by 0.3- to 18.4-fold, except one specimen that exhibited 24,000-fold higher infectivity than the specimen not treated with trypsin (Fig. 3B). Thus, trypsin treatment increased the infectivity of the omicron variant to a lower extent than that of the delta variant. We observed no significant difference in the Ct values of specimens with or without increased infectivity. We speculate that the effect of trypsin treatment on increasing infectivity is not simply affected by viral load.

**Fig. 3.**
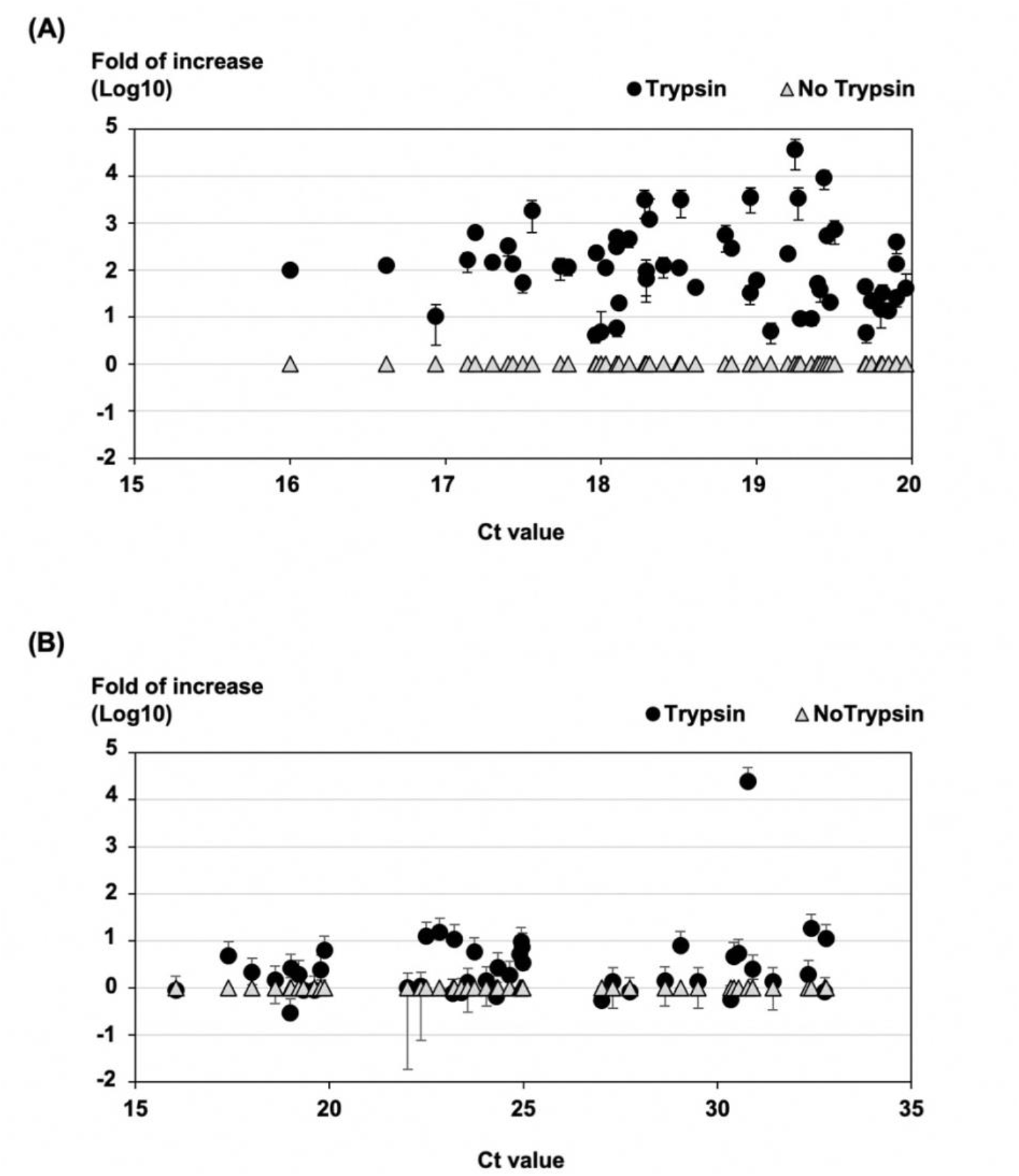
Effect of trypsin treatment on SARS-CoV-2 infection using clinical specimens. Comparison of clinical specimens of the (A) delta (n = 56) and (B) omicron (n = 44) variants by Ct value at PCR testing. The load of viral genome after viral amplification with trypsin treatment is expressed as the infection enhancement ratio, where the amount of viral genome after viral amplification without trypsin treatment is set as 1. ●; Relative viral genome quantity of trypsin (400 μg/mL)-treated specimens, △; Relative viral genome quantity of specimens not treated with trypsin.

### Enhancement of SARS-CoV-2 infection by culture supernatants of anaerobic bacteria

We examined whether culture supernatants of anaerobic bacteria have the same effect as trypsin on enhancing infection (Fig. 4A–D). Six clinical COVID-19 specimens, including the SARS-CoV-2 delta variant, were used to determine whether culture supernatants of oral anaerobic bacteria (*Prevotella melaninogenica, Prevotella intermedia*, and *Fusobacterium necrophorum*) increased infectivity. Treatment with culture supernatants of *P. melaninogenica* or *P. intermedia* increased infectivity negligibly. By contrast, treatment with culture supernatant of *F. necrophorum* enhanced viral infection by ∼10-fold compared to treatment with medium alone.

**Fig. 4.**
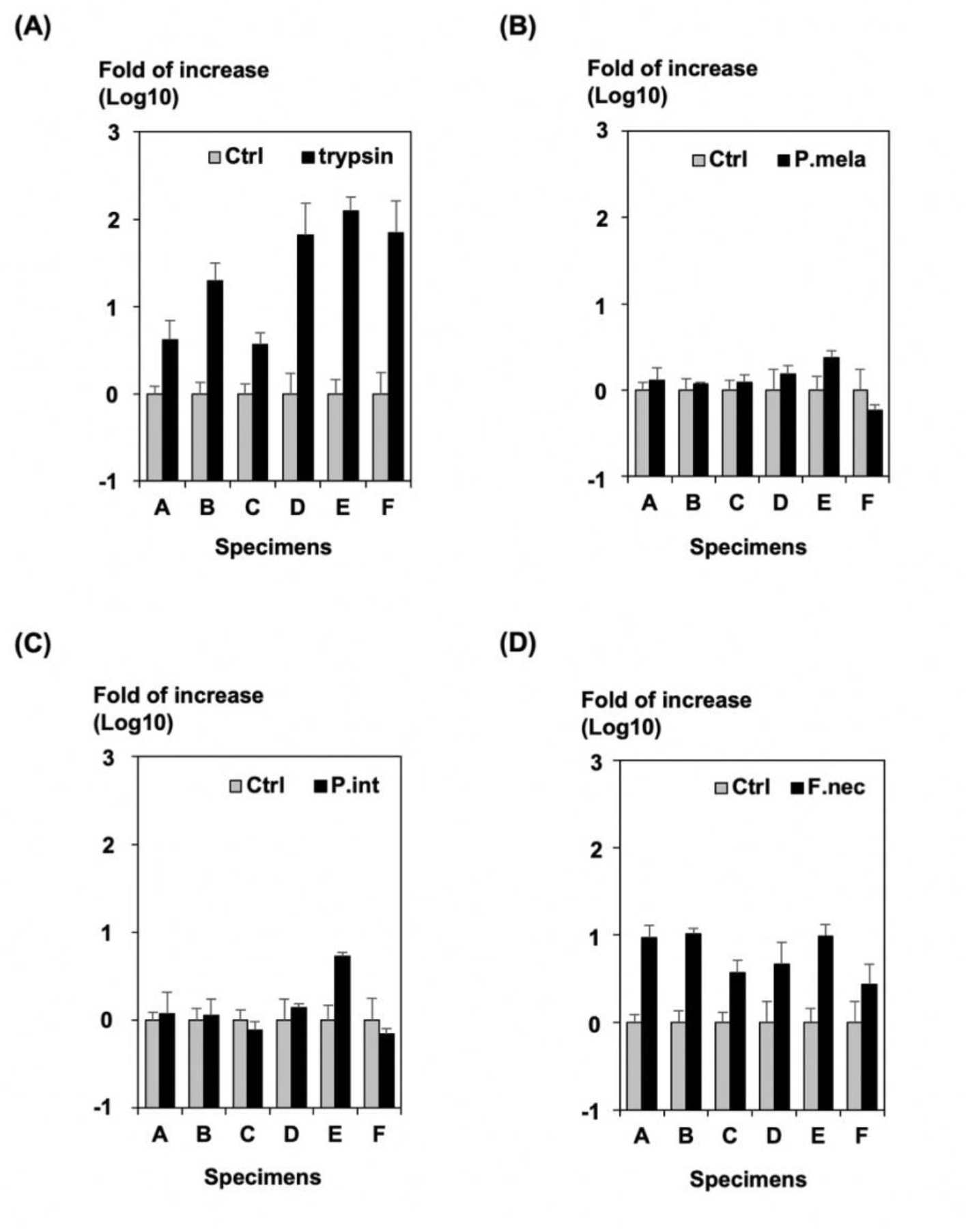
Effect of anaerobic oral bacterial culture supernatant on the infectivity of SARS-CoV-2 using clinical specimens. After adding the 6 clinical specimens (A–F) containing the SARS-CoV-2 delta variant to the cells, the increase in infectivity of viruses was calculated after treatment with (A) trypsin (800 µg/mL) and culture supernatants of (B) *Prevotella melaninogenica*, (C) *Prevotella intermedia*, and (D) *Fusobacterium necrophorum* based on the infectivity of the virus treated in the control medium. The results shown are from three independent assays, with error bars representing standard deviations.

## Discussion

Trypsin treatment of cells increases viral infectivity and increases membrane fusion in animal and human coronaviruses, including SARS-CoV (4, 6, 7). SARS-CoV-2 virus entry was reported to increase in the presence of trypsin (2, 8). Our findings revealed that trypsin treatment of cells increased SARS-CoV-2pv infectivity as well as infectivity of SARS-CoV-2 present in clinical specimens derived from the nasal and oral cavities of patients with COVID-19. The increase in infectivity was different among SARS-CoV-2 variants. In clinical specimens from patients with COVID-19, the delta variant showed up to tens of thousands-fold increase in infectivity after trypsin treatment, whereas the omicron variant showed only a few to several dozen-fold enhancement. We observed no significant difference in Ct values between clinical specimens in which infectivity increased greatly after trypsin treatment and those in which infectivity did not increase, suggesting that the increase in infectivity after trypsin treatment is not simply affected by viral load. Although this difference is thought to be caused by differences in the components contained in the clinical specimens, the detailed reason is unknown and further investigation will be needed.

Studies have shown that infection by members of the *Orthomyxoviridae* and *Paramyxoviridae* families, including the influenza virus, are facilitated by some proteases (9-12). Because the envelope glycoproteins of these viruses are not completely cleaved in *de novo* synthesized cells, the viral particles produced from these cells contain partially-cleaved or uncleaved glycoproteins. Treatment with proteases cleaves the glycoproteins on the virions, thus enhancing their infectivity. Thus, trypsin acts directly on virions to increase infectivity. However, the infectivity of SARS-CoV was reported to increased when S protein is cleaved by proteases after binding to the ACE2 receptor (4). In SARS-CoV-2 infection, the increase in infectivity after treatment with trypsin or other proteases may be due to a mechanism different from that for influenza virus or other viruses. Each SARS-CoV-2pv variant was pretreated with trypsin before being used to infect the cells; however, this treatment did not increase the infectivity of SARS-CoV-2pv. In other words, the protease can induce the fusion activity of S protein only after the virus binds to the ACE2 receptor. Studies have shown that the omicron variant uses a TMPRSS2-independent entry pathway (13, 14). In our study, treatment with trypsin did not significantly increase the infectivity of the omicron variant, suggesting that trypsin treatment enhances TMPRSS2-mediated cell entry.

In SARS-CoV-2 infection, syncytium formation in infected cells has been reported to play a role in viral replication and pathogenesis of severe COVID-19 (15, 16). Moreover, cellular membrane fusion activity of the omicron variant was reported to be lower than that of the delta variant (13). Studies have reported that cell membrane fusion is activated by treatment with buffers with an acidic pH (17). In this study, we found that trypsin-like proteases activate cell membrane fusion.

Furthermore, we found that the infectivity of SARS-CoV-2 from the nasal cavity increased in the presence of culture supernatants of oral anaerobic bacteria, such as *F. necrophorum*. It is unclear why SARS-CoV-2 infectivity increased only when culture supernatants of *F. necrophorum* were used and not *P. melaninogenica* and *P. intermedia*. Therefore, the amount and type of proteases in the culture supernatant must be evaluated. These are a type of commensal bacteria that are present in the oral cavities of healthy people and have been reported to be involved in the exacerbation of periodontal disease and gingivitis (18, 19). Studies have shown differences in the nasopharyngeal microbiome of patients with COVID-19 (20). In addition, although operational taxonomic units classified as *Prevotella* were found to be significantly more abundant in patients who developed more severe COVID-19, the mechanism of interaction between the salivary microbiome and SARS-CoV-2 remains to be elucidated (21).

In summary, our findings revealed that SARS-CoV-2 in the nasal or oral cavity of patients with COVID-19 is activated by the action of proteases. Therefore, keeping the oral cavity clean on a daily basis and reducing components derived from oral indigenous bacteria will help prevent SARS-CoV-2 infection.

## Materials and Methods

### Specimens

The clinical specimens used in this study were SARS-CoV-2-positive nasal and nasopharyngeal swabs collected in August 2021 to March 2022 from patients with COVID-19 at the Toyama Institute of Health. Specimens identified as delta variants were suspected delta variants in the mutation screening PCR test (L452R), and specimens identified as omicron variants were identified as BA.1 variants by genome analysis using next-generation sequencing. Ct values were obtained from PCR tests conducted as administrative tests at the Toyama Institute of Health using the SARS-CoV-2 direct detection RT-qPCR test kit from Takara Bio Inc. (Shiga, Japan). The study was approved by the Ethics Review Committee of the Toyama Institute of Health (R2-12).

### Plasmids, cells, and viruses

The expression plasmids pCAGGs-pm3-SARS2-S-Hu_d19_D614G, pCAGGs-pm3-SARS2-S-Hu_d19_B.1.1.7, pCAGGs-pm3-SARS2-S-Hu_d19_B.1.617.2, and pCAGGs-pm3-SARS2-S-Hu_d19_BA.1, which encode S genes of Wuhan, alpha, delta, and omicron BA.1 variants, respectively, were kindly provided by Drs. C. Ono and Y. Matsuura, Research Institute for Microbial Diseases, Osaka University. The expression plasmid pCAGGS-eGFP, which encodes enhanced green fluorescent protein (eGFP), was constructed as described (22). VeroE6 and 293T cells were obtained from the American Type Culture Collection (Summit Pharmaceuticals International, Japan) and cultured in Dulbecco’s modified Eagle’s medium (DMEM; Sigma-Aldrich, St. Louis, MO), containing 10% heat-inactivated fetal bovine serum (FBS).

Pseudotyped VSV bearing S protein of SARS-CoV-2 was generated as described (23). Briefly, 293T cells were transfected with the expression plasmids. After 24 h of incubation, the transfected cells were infected with G-complemented (*G) VSVΔG/Luc (*G-VSVΔG/Luc) at a multiplicity of infection of 0.5. After the cells adsorbed the virus, they were washed four times with DMEM containing 10% FBS. After 24 h of incubation, culture supernatants containing pseudotyped VSVs were centrifuged to remove cell debris and stored at −80°C until use.

### Real-Time PCR

VeroE6 cells seeded in 96-well culture plates were treated with DMEM containing 8% FBS on ice for 10 min. Approximately 5 μL centrifugal supernatant of nasal and nasopharyngeal swabs, which were positive for SARS-CoV-2 in RT-qPCR, was added to the cells. After adsorption on ice for 60 min was complete, supernatants including the swabs were removed, and the cells were treated for 5 min with the indicated concentrations of trypsin or bacterial culture supernatants in DMEM containing 2% FBS that was prewarmed at room temperature. Then trypsin or bacterial culture supernatant was removed, and cells were cultured in DMEM containing 8% FBS at 37°C for 24 h. RT-qPCR was performed to estimate the amounts of RNA. For RT-qPCR testing, we used the QIAamp Viral RNA Mini Kit (QIAGEN, Cat. No. 52904, Germantown, MD), which targets the N gene of SARS-CoV-2, and the assays were performed using QuantStudio 5 (Thermo Fisher Scientific Inc. Waltham, MA) (24). RT-qPCR analysis was performed under the following conditions (reverse transcription: 50°C for 5 min, 95°C for 20 s; PCR: 95°C for 75 s and 45 cycles of 95°C for 15 s, 60°C for 60 s). To quantify virus titer produced in the cells, we collected the cell culture supernatants, and determined the amount of RNA diluted 10-fold using RT-qPCR.

### Trypsinization experiments for SARS-CoV-2pv

To examine the effect of trypsin after viral infection, VeroE6 cells seeded in 96-well culture plates were treated with DMEM containing 10% FBS on ice for 10 min. Approximately 100 μL of SARS-CoV-2pv (Wuhan, alpha, delta, and omicron) or VSVpv were added to the cells. After viral adsorption on ice for 60 min, the virus was removed, and the infected cells were treated for 5 min with various concentrations of trypsin (Sigma, T1426) in DMEM that was prewarmed at room temperature. After trypsin solution was removed, cells were cultured in DMEM containing 10% FBS at 37°C for 24 h.

To examine the effect of trypsin on viral particles, SARS-CoV-2pv or VSVpv was exposed to various concentrations of trypsin in DMEM for 5 min at 37°C. To inactivate trypsin, FBS was added at a final concentration of 20%. Then 20 μL of sample was added to each well, and the cells were incubated at 37°C for 24 h. The infectivity of both SARS-CoV-2pv and VSVpv was assessed separately by measuring luciferase activity. The relative light unit value of luciferase was determined using the PicaGene Luminescence Kit (TOYO B-Net Co., Ltd., Tokyo, Japan) and GloMax Navigator System G2000 (Promega Corporation, Madison, WI), according to the manufacturer’s protocol.

### Syncytium formation

To examine syncytium formation using S protein of SARS-CoV-2 variants after trypsin treatment, VeroE6 or 293T cells were transfected with the indicated S protein of SARS-CoV-2- or VSVG-expressing plasmid together with pCAGGS-eGFP. At 48 h post-transfection, the cells were treated for 5 min with 800 µg/mL trypsin in DMEM containing 2% FBS that was prewarmed at room temperature. The trypsin-containing media was replaced with DMEM containing 10% FBS and incubated for 24 h. The cell monolayers were then observed for syncytium formation under a fluorescence microscope.

### Addition of anaerobic bacterial culture supernatant to clinical specimens

We examined whether the culture supernatant of oral anaerobic bacteria increased SARS-CoV-2 infectivity. Three species of oral anaerobic bacteria: *P. melaninogenica* JCM6325, *P. intermedia* JCM11150, and *F. necrophorum* JCM3718 were obtained from Japan Collection of Microorganisms (RIKEN BRC, Ibaraki, Japan) and cultured using Brain Heart Infusion medium (Becton, Dickinson and Company, Cat. No. 237500, Franklin Lakes, NJ) supplemented with hemin (Sigma-Aldrich, H9039), menadione (Sigma-Aldrich, M9429), L-cysteine (Sigma-Aldrich, Cat. No. 168149), and yeast extract (Thermo Fisher Scientific Inc. Cat. No. 212750). The bacteria were cultured in an anaerobic environment for 72 h and filtered through a 0.22-µm filter. Six clinical specimens of the delta variant, which showed high increase in infectivity after trypsin treatment, were added to VeroE6 cells that had been chilled on ice for 10 min. Incubation was performed for 60 min on ice for the virus to adsorb on the cells. Then the virus was removed, and the cells were treated with the culture supernatant of each anaerobic bacteria at 37°C for 5 min. Then the culture supernatant was removed, and the cells were cultured in DMEM containing 10% FBS at 37°C for 24 h. To measure the viral titer produced in the cells, the cell culture supernatants were collected, and the amount of RNA diluted 10-fold was determined by RT-qPCR.

## Author Contribution

Conceptualization and methodology, E.I., Y.M., K.O. and H.T.; sample collection, E.I., T.S., S.Y., Y.S., M.I., N.I., S.H.; investigation. E.I., T.S., S.Y., Y.M., and H.T.; writing–original draft presentation, E.I., Y.M., and H.T.; writing–review and editing, E.I., Y.M., K.O., and H.T.; supervision, K.O., and H.T.; funding acquisition, Y.M. and H.T. All authors have read and agreed to the published version of the manuscript.

## Funding

This work was supported in part by a grant-in-aid from the Japan Agency for Medical Research and Development (AMED) (Grant No. JP20he0622035, JP21fk0108588 and JP22fk0108133) and in part by a grant-in-aid from the Tamura Science and Technology Foundation.

## Acknowledgments

Tragically, Sumiyo Hasegawa passed away on 23 May 2022. Her memory will remain among all of us who shared part of our lives and enjoyed her company. Because of her enthusiasm, dedication, and motivation, she will be remembered as a wonderful person. We sincerely would like to thank Yoko Kanamori and Izumi Kawaguchi for technical and secretarial assistance.

## Conflict of interest

The authors declare no conflict of interest.

## Notes

### Competing Interest Statement

The authors have declared no competing interest.

